# Asymmetric regeneration dynamics in zebrafish pectoral fins

**DOI:** 10.64898/2026.10.26.761995

**Authors:** Rocky Diegmiller, Sushant Bangru, Kelsey Oonk, Kenneth D. Poss, Stefano Di Talia

## Abstract

Appendage amputation in regeneration-capable systems results in the full re-formation of lost tissue and cell populations. Despite an appreciation of this phenomenon in a select number of organisms, there is still much to quantitatively understand about the tissue growth dynamics during limb regeneration. To address these gaps in knowledge, we developed novel methods that enable longitudinal, quantitative imaging of zebrafish pectoral fin regeneration and uncovered a persistent spatial asymmetry in growth dynamics between anterior and posterior regions of the regenerating fin. Spatiotemporally resolved transcriptomics suggests differential anteroposterior regulation of the Mitogen-activated protein kinase (MAPK) and Wnt signaling pathways in pectoral fins. This asymmetry was also detected using quantitative imaging of biosensors reporting canonical Wnt and Erk pathway activity. In addition to known static spatially asymmetric genes, bulk transcriptomic analysis uncovered a large subset of genes with temporally asymmetric peaks in expression levels between anterior and posterior regions of the fin across regeneration timepoints, suggesting a global differential regulation of the regenerative program. Taken together, our results establish pectoral fins as a platform for quantitative analyses of appendage regeneration, and they identify persistent spatial asymmetries in growth, gene expression, and signaling activity during regeneration.

---

In contrast to mammals, some vertebrates, like salamanders and bony fish, can regrow damaged organs and appendages back to their original size and shape^1–5^. Many genes and signaling pathways involved in the regenerative process also underlie the uncontrolled proliferation observed in tumor growth. In a regeneration context, however, these signals direct the program to grow back only what was lost^6,7^. Although this phenomenon – termed positional memory – is well-appreciated, the growth dynamics and underlying signals promoting proliferation and movement of cell populations throughout regeneration require additional resolution^1,8–11^. Therefore, the study of regenerating tissues with intrinsic asymmetries and gradients can provide a powerful framework to better analyze how tissue re-forms after injury.

Zebrafish pectoral fins are attractive models to quantitatively study regeneration. These fins exist as a pair of appendages located just posterior to the gills and are the evolutionary precursor of vertebrate forelimbs^12–14^. In contrast to the more commonly studied caudal fin, pectoral fins also possesses clear morphological gradients in fin ray size from anterior (thicker, longer) to posterior (thinner, shorter), as well as gene expression gradients along the anteroposterior (A/P) axis, as in vertebrate limbs^9,15^. Zebrafish pectoral fins are also transparent and thin, enabling spatiotemporally resolved quantification of gene expression and cellular dynamics at high resolution^15^.

Structurally, each pectoral fin is enveloped by a surface layer of epithelial cells and contains 11-13 bony fin rays, while each ray is made up of two osteoblast hemirays filled with fibroblasts and interconnected via mesenchymal, fibroblast-rich tissue^12,15^. Upon amputation, osteoblasts in the fin rays dedifferentiate and fibroblasts collect at the base of the amputation site to form populations of cells known as blastemas. During the first days after amputation, fibroblasts begin to proliferate and osteoblasts re-differentiate, starting the roughly four-week regrowth process^3,16,17^. Due to its proximity to the gills, previous work describing pectoral fin regeneration removed the entire appendage from the fish when imaging, preventing longitudinal studies^15,18,19^. To better answer the questions of signaling and morphological dynamics during limb regeneration, we have developed longitudinal imaging methods for pectoral fins.

## Results

### Anteroposterior heterogeneity of pectoral fin ray regeneration dynamics

After amputation, respondent cell populations begin to undergo changes in their gene expression profiles, as well as their proliferative and migratory capacities to begin the wound healing and outgrowth process to reform lost tissue^3,16,17^. Previous work has shown that pectoral fins, like caudal fins, will fully regenerate their fin rays within 4 weeks post amputation. Unlike caudal fins, however, a highly-resolved picture of pectoral fin regeneration dynamics across the fin has not been fully revealed, as techniques for the longitudinal imaging of the same animal had not been established^15,18,19^. To address this, we designed a 3D-printable mold consisting of a partial circular base with a small, sloped region removed from the center, along with a small triangular prism-like piece that attaches to form an adjustable canal (Fig. S1). After applying anesthesia, fish can be inserted ventral side up and held firmly in place while its pectoral fin is splayed out against the base. Once imaged, the fish can be easily removed and recovered for further acquisitions.

With this device, we performed amputations and measured the regeneration dynamics of individual fin rays (Fig. 1A). Throughout, as is true for all experiments performed in this study, we focused only on female zebrafish pectoral fins, as previous works have demonstrated the tendency for male pectoral fins to exhibit regeneration defects^18,19^. Using iridectomy scissors, a physical amputation was performed on *osx:H2A-mCherry* animals to remove a large portion of the pectoral fin^15,20^. This proximal amputation, which removed 60% of anterior rays and 45% in posterior rays, was performed on 10 fish, which were periodically mounted and imaged over 33 days (Fig. 1B). Initially, we observed at early times (2 days post amputation, dpa), little outgrowth was present, consistent with the blastema formation stage of regeneration. Interestingly, however, the following collected timepoint at 5 dpa revealed that posterior fin rays had regrown significantly more tissue relative to length amputated than anterior rays (33 ± 2%, SEM and 35 ± 2%, SEM of rays 6 and 7, respectively, compared to 19 ± 2%, SEM and 22 ± 2%, SD of rays 2 and 3, respectively). This initial early growth spike in posterior fin rays, however, waned over time, as anterior ray regeneration rates eventually caught up such that all rays reached their unamputated lengths at roughly the same time (Fig. 1C).

**Figure 1:**
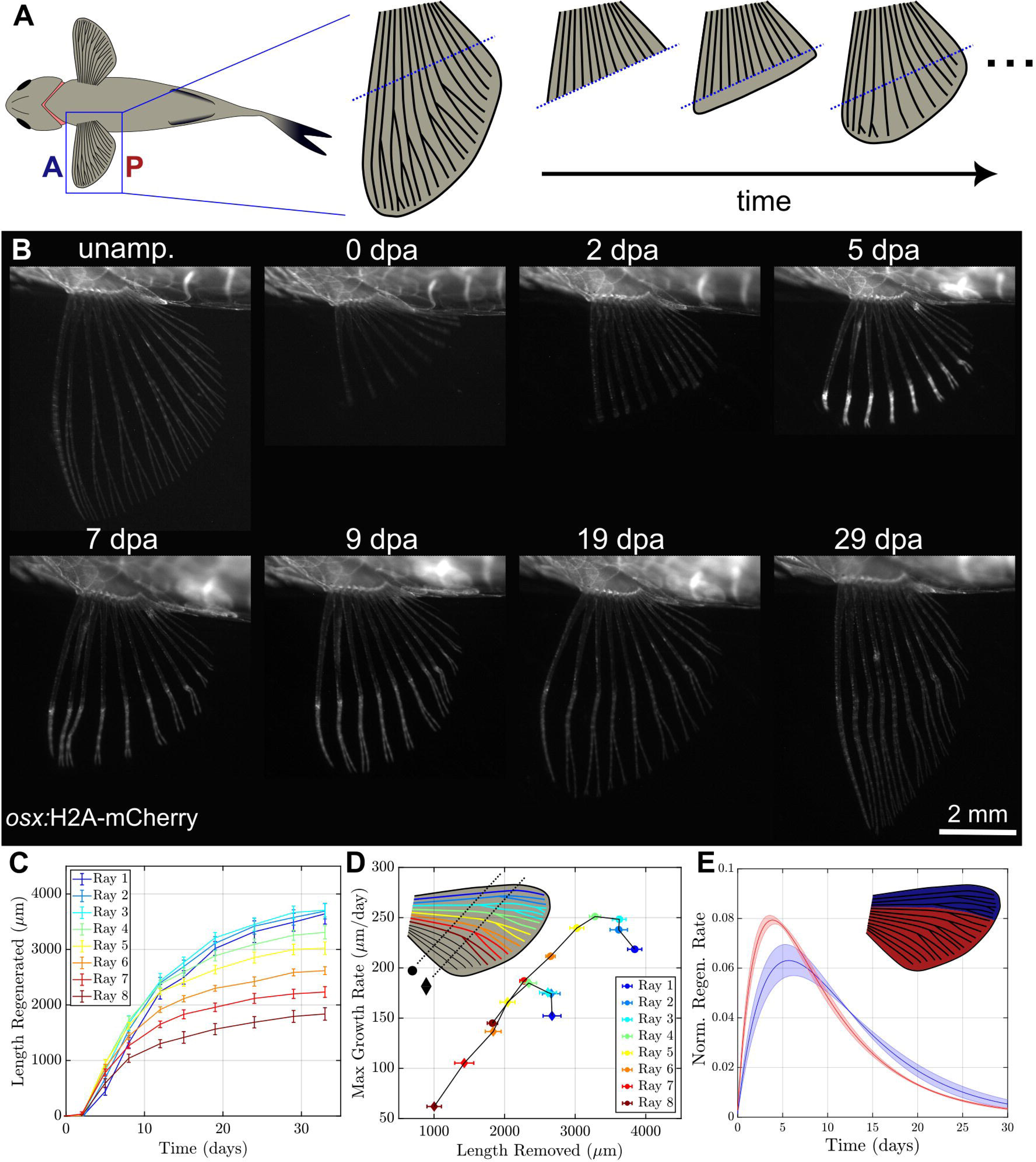
Longitudinal dynamics of female pectoral fins reveals spatially distinct growth dynamics. A) Left: schematic of zebrafish placed upside-down showing paired pectoral fin locations and size gradient of individual fin rays from anterior (A) to posterior (P). Right: Schematic of zebrafish pectoral fin regeneration dynamics. B) Example whole-fin, stereoscopic images of regenerating zebrafish pectoral fin expressing *osx:H2A-mCherry* at unamputated, 0, 2, 5, 7, 9, 19, and 29 dpa timepoints highlight dynamics of regenerating fin rays, from initial outgrowth to end-stage regeneration (scale bar = 2 mm). C) Plot of average total length regenerated as a function of time (n=10) for the distal amputation (∼50% removal) experiment. Rays 1 (most anterior, dark blue) through 8 (most posterior, dark red). Across fish, the most anterior rays’ dynamics (1-3) appeared qualitatively distinct from the posterior rays’ (4-8), prompting further analyses (error bar = SEM). D) Estimated maximum growth rate of each ray (in μm/day) as a function of average fin length removed (in μm) has a different relationship between anterior (1-3) and posterior rays (4-8). This distinction was not a function of amputated length, as both the distal (circle, n=10) and proximal amputation (diamond, n=11) experiments showed the same trend (error bar = SEM). E) Plot of fraction regenerated over time (per day) as a function of time. After grouping the fin into anterior (n=3) and posterior (n=5) portions, the average normalized growth rate over time showed distinct dynamics between anterior and posterior rays (error bar = SD).

This experiment was repeated, this time with a less severe amputation on 11 fish, where around 40% of anterior rays down to 25% of posterior rays were removed. Despite this decrease in amputated lengths, the same pattern of early asymmetry was observed again, with the anterior rays again catching up by the end of regeneration (Fig. S2). To account for larger tissue losses, we additionally performed amputations of longfin (*lof*) fish, whose pectoral fin rays are substantially and proportionally longer compared to wildtype^21^. After removing roughly 50-60% of these elongated rays across the fin, we again found the same pattern of regeneration rate differences within this system (Fig. S2). Consistently, we observed posterior fin rays growing proportionally more tissue following amputation, while anterior rays grew back initially more slowly, but sustained more growth over a longer time. Across experiments, these observed growth differences suggested the observed phenomenon was not simply due to a difference in length amputated.

To better understand this behavior and fully define the distinct anteroposterior regions with respect to growth, we further quantified and generated a minimal model to describe the spatiotemporal asymmetries of regenerating pectoral fin rays. For each experiment, fin ray trajectories were collected based on A/P identity and fitted with a bi-exponential curve to estimate growth rates. This choice of fitting was made in part due to the qualitative logistic-like growth trajectories observed for each fin ray, which resemble an out-of-equilibrium, overdamped second order system as it relaxes to a set point – in this case, as the regenerating fin ray grows back its missing length^22^.

By comparing estimated maximum growth rate from these fits as a function of average length amputated, a strong linear relationship was observed for posterior fin rays, but this relationship did not hold for the three most anterior rays in the fin (Fig. 1D). Similarly, there was a consistent difference across experiments between anterior and posterior rays in their estimated time to reach maximum growth rate: posterior rays reached this maximum around 4 dpa, while the anterior rays peaked around 6 dpa (Fig. 1E). With these distinct regions, along with the known gene expression differences present in the pectoral fins, we sought to identify genes whose expression could correlate with the observed growth rate differences between regions of the regenerating fin.

### Cell type and signal pathway enrichment of growth rate and gene expression dynamics

Previous work studying pectoral fin regeneration performed bulk RNA-seq analysis only on unamputated anterior and posterior fin rays to identify spatially asymmetric genes in unperturbed conditions^15^. In addition, qPCR was also performed on 4 dpa fins and compared anterior, medial, and posterior sections^15^. However, no analyses have been devoted to analyzing gene expression dynamics of regenerating pectoral fins over time. Therefore, we first wanted to classify cell types and genetic spatiotemporal heterogeneity present across the fin to identify potential distinctions during early regeneration that could contribute to the observed growth rate asymmetry by performing single-nucleus RNA sequencing (snRNA-seq) of adult female zebrafish pectoral fins. To do this, we amputated ∼50% of each pectoral fin from around 75 fish. Each fin was then sectioned into fin rays 1 and 2 (AP1), 3 and 4 (AP2), 5 and 6 (AP3), 7 and 8 (AP4), and the most posterior remainder (AP5). At 2, 4, and 7 dpa, each of 25 female fish were re-amputated and sectioned into AP1-AP5 to collect from these regions (Fig. 2A).

**Figure 2:**
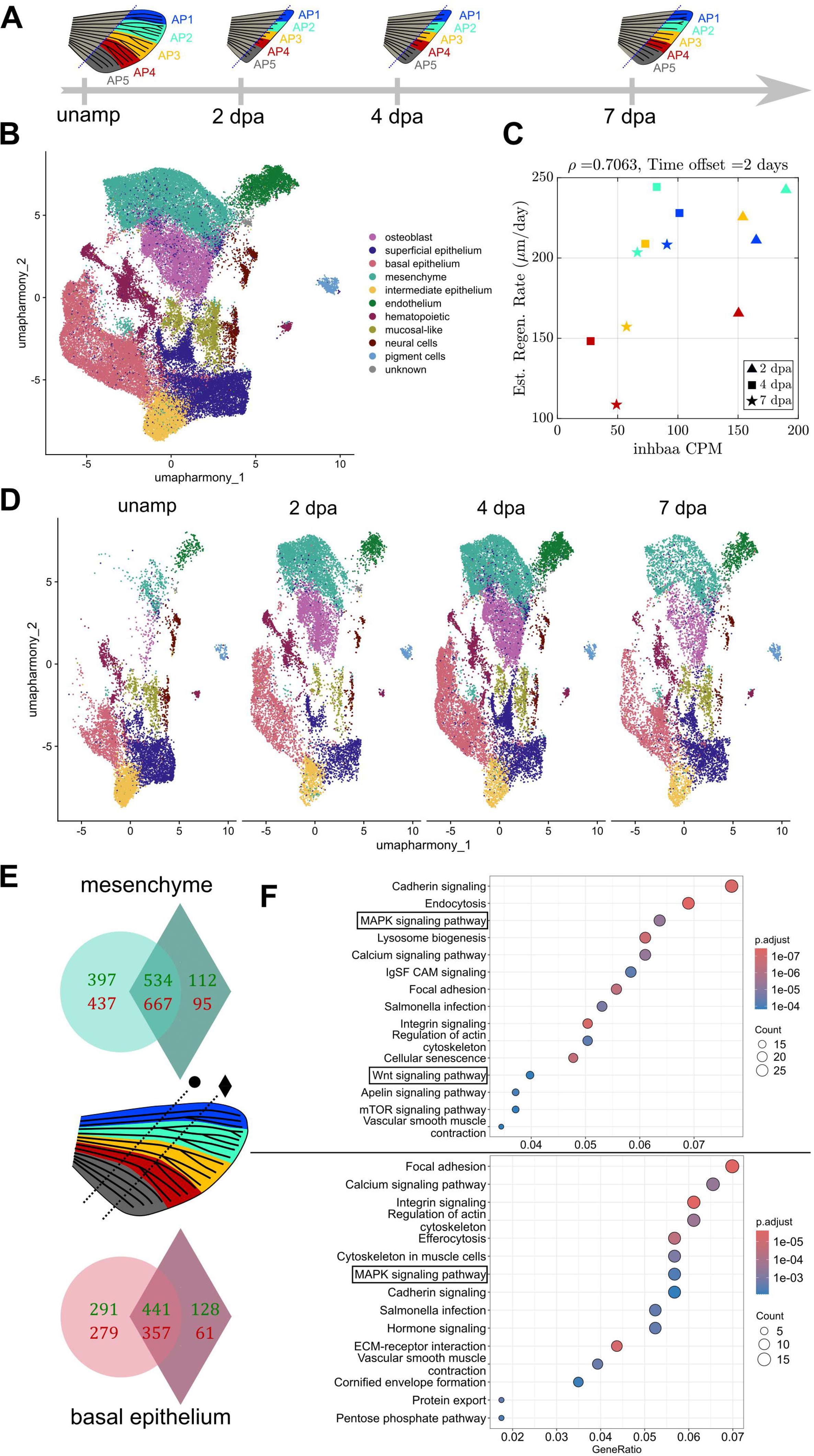
snRNA-seq analyses highlight regenerative cell types and enriched pathways. A) Timeline of single-nuclei tissue amputation and collection, splitting into AP1-5 throughout regeneration timepoints 2, 4, and 7 dpa. B) UMAP representation of cells collected from experiment after quality control (n=50,784) with cell type labels. AP1-5 across the four collected timepoints generated 19 clusters that were binned into 15 cell types based on known cell type markers and differentially expressed genes within each cluster. C) Binned estimated growth rate from the distal amputation experiment with added time offset as a function of gene CPM across 2, 4, and 7 dpa for AP1-4 for both basal epithelium and mesenchymal cell types generated a Spearman correlation, ρ, for all offsets between 0 and 4 days. The maximum (in absolute value) Spearman correlation was kept for all 26,395 genes collected from the snRNA-seq, with |ρ| > 0.7 being used as a cutoff to show a strong relationship between growth rate and gene expression. D) UMAP plots across cell type for each collected regeneration timepoint (2, 4, and 7 dpa) shows a large and consistent proportion of mesenchymal (28.3%, 26.3%, and 19.6%, respectively) and basal epithelial cells (17.5%, 20.5%, 19.4%, respectively), making up a plurality of all cells collected. E) For each of the mesenchymal (top) and basal epithelial (bottom) cells, those genes with a sufficient positive or negative Spearman correlation in both the proximal and distal amputation experiments were kept, yielding 1201 and 798 genes from analyzing expression from mesenchymal and basal epithelial cells. F) KEGG pathway analysis on genes from the mesenchymal (top) genes revealed MAPK and Wnt signaling pathways were enriched, prompting further analyses into how these pathways may be involved in guiding regeneration rates in pectoral fins. Similarly, the MAPK pathway was also enriched from analyzing basal epithelial genes. Overall, there was an additional notable enrichment of pathways involving cell mechanics and ion signaling.

After initial quality control filtering, 50,784 nuclei were retained for further analyses. Unsupervised clustering generated 19 distinct clusters of cells based on transcriptional profiles, independent of spatiotemporal identity and cell type identities were assigned based on the presence of previously characterized marker genes, identifying populations of basal epithelial, endothelial, hematopoietic, intermediate epithelial, mesenchymal, mucosal-like, osteoblast, neural, pigment, and superficial epithelial cells^23–27^ (Fig. 2B). Across timepoints and spatial strata, we found basal epithelial cells (9,838, 19.4%) and mesenchymal cells (11,175, 22.0%) to be the most highly enriched (Fig. 2D). As these specific cell types have also been previously shown to be involved in signaling during appendage regeneration, we used the gene expression information from these cell types to perform downstream analyses^11,28,29^. Within these populations, we searched for genes with expression levels correlated with the average growth rates for the regions AP1, AP2, AP3, and AP4 throughout regeneration derived from the previously collected and fit growth data.

In parallel, for each of the 26,395 genes identified, pseudobulk counts per million (CPM) for each cell type were generated for each collected timepoint and spatial region. These values were plotted against estimated growth rates, first at 2, 4, and 7 dpa, in addition to allowing for an offset (up to 4 dpa) to account for possible time delay of integrating signal and lag effect on growth (Fig. 2C). Across time delays, we selected only those genes with a Spearman correlation coefficient > 0.7 or < −0.7 and CPM above 1 over some collected timepoints, yielding 931 positively correlated and 1104 negatively correlated mesenchymal genes from the proximal amputation growth rates, along with 646 positively correlated and 762 negatively correlated basal epithelial genes (Fig. 2E). Performing the same analysis with the distal amputation data generated 732 positive correlated and 636 negatively correlated mesenchymal genes, along with 569 positively correlated and 418 negatively correlated basal epithelial genes, of which a total of 1201 mesenchymal and 798 basal epithelial-driven genes were found in both cases and used for further downstream analyses (Fig. 2E).

Using the Kyoto Encyclopedia of Genes and Genomes (KEGG), pathway analysis was performed on these lists of overlapping mesenchymal and basal epithelial genes^30^. Notably, we found the Mitogen-activated protein kinase (MAPK) signaling pathway to be highly enriched in both basal epithelial and mesenchymal cells, along with calcium signaling, Wnt signaling and mTOR signaling also upregulated in mesenchymal cells (Fig. 2F). Notably in both cell types, there was significant upregulation of mechanical signaling pathways, like cadherin signaling, focal adhesion, integrin signaling, and actin regulation. However, based on previous works in the caudal fin that have demonstrated the importance and timing of MAPK and Wnt signaling during appendage regeneration, we focused on these pathways for further analyses^11,29,31–34^.

### Longitudinal imaging of signal pathway dynamics reveals spatiotemporal asymmetries

Previous work has shown that the activity of Erk, a downstream component of the MAPK pathway, plays an important role in appendage regeneration through guiding osteoblast proliferation^11,33^. In addition, previous work has shown Wnt signaling is involved in distal mesenchymal cells adopting a pre-osteoblast fate via *runx2*, leading to osteoblast maturation during regeneration^35,36^. Based on these findings and the results of our pathway analysis from the snRNA-seq studies, we asked what role Erk and Wnt signaling might play and if there may be dynamic asymmetries that align with our observed growth differences throughout the fin.

Quantifying activity dynamics, however, required longitudinal imaging of pectoral fin rays at the confocal level to measure activity within individual cells. To address this, we utilized a modified dissection scope mold made of clear resin that was placed within a plexiglass dish, in a similar manner to previous works performing longitudinal imaging of regeneration^11,20,33^. Anesthetized fish were placed upside down and 2-3 drops of cooling 1% agarose placed on their pectoral fin and submerged in diluted anesthetic solution for the duration of imaging. To minimize the amount of time the fish stayed under deep anesthesia before being revived, only one anterior (ray 2) and posterior (ray 6 or 7) fin ray were imaged throughout these studies.

As recent work has characterized Erk activity in osteoblasts as a readout for regeneration progression, we sought to analyze the dynamics of this sensor, also in osteoblasts, to examine if there may be emergent regional differences in Erk activity over time^11^. Thus, to analyze Erk dynamics in osteoblasts, we imaged fish containing *osx:H2A-mCherry* and *osx:ErkKTR-mCerulean* transgenes, which generates a readout of Erk activity based on the ratio of cytoplasmic to nuclear-localized fluorescence of the sensor^11,33^ (Fig. 3A). Across all imaged rays, longitudinal imaging of this reporter across 6 fish ranging from 3 to 11 dpa showed average Erk activity is inversely related to fraction regenerated of fin ray length, similar to previous results in the caudal fin^11^ (Fig. 3B). In comparing the average Erk activity levels between fin regions, we observed slightly overall lower levels in posterior rays (0.20 ± 0.02, SEM) compared to the anterior (0.25 ± 0.03, SEM) at 9 dpa, when the posterior rays had grown proportionally more of their amputated lengths (0.41 ± 0.03, SEM) than their anterior counterparts (0.39 ± 0.01, SEM), providing evidence of growth-driven asymmetry in these regions (Fig. 3C). Across fish, we observed a trend of lower average Erk activity between anterior and posterior rays over time, although due to the constraints of the imaging process preventing respiration, we were unable to image dynamics of these fins beyond 11 dpa.

**Figure 3:**
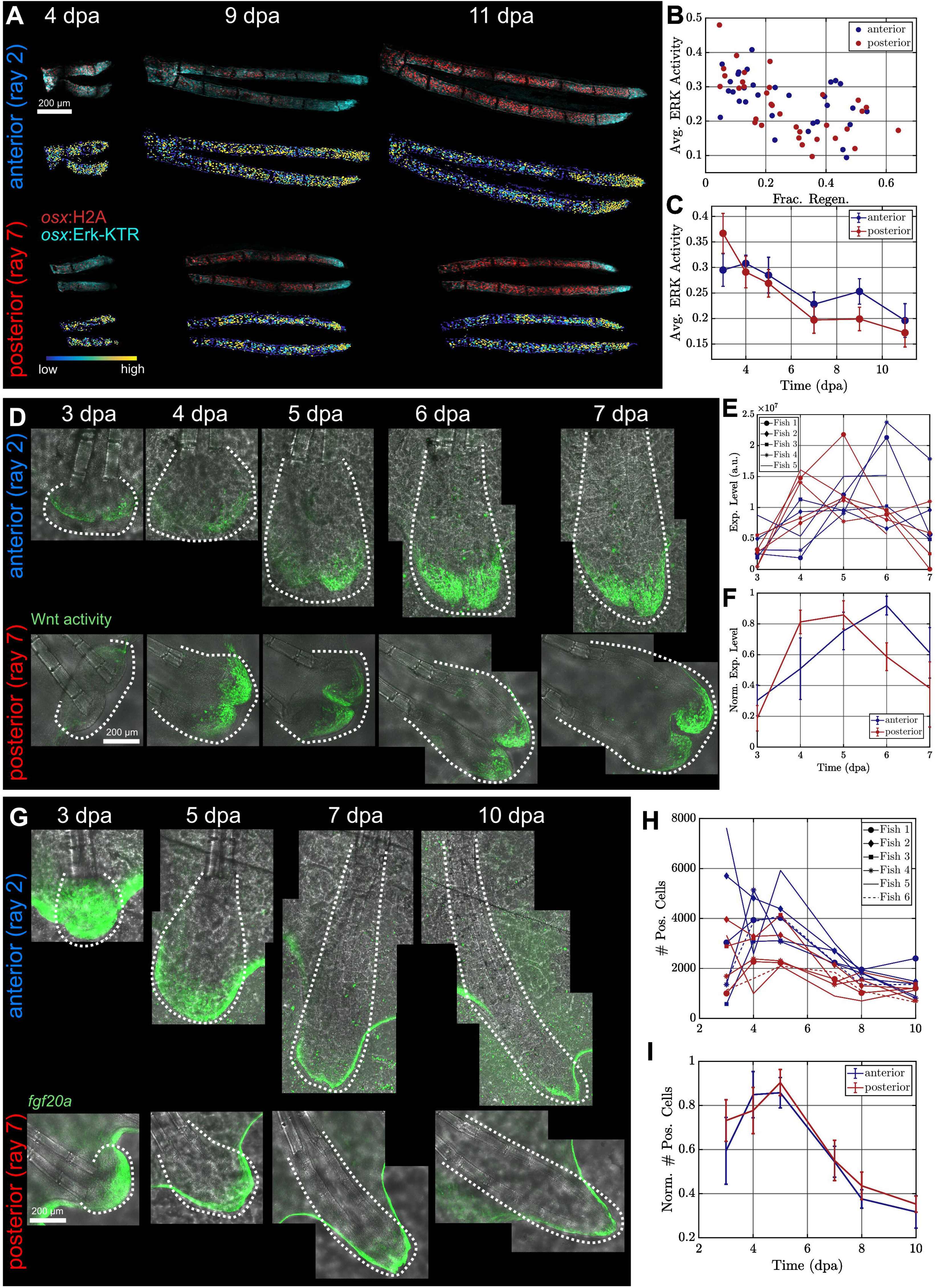
Longitudinal confocal microscopy shows anteroposterior asymmetry in MAPK and Wnt pathway activity dynamics, but this asymmetry is not associated with *fgf20a* expression. A) Maximum projection images of representative anterior and posterior rays at 4, 9, and 11 dpa co-expressing *osx:H2A-mCherry* and *osx:ErkKTR-mCerulean*, along with the processing activity levels for each osteoblast cell in the regenerating fin ray (blue = low, yellow = high). B) Average ERK activity level appears inversely related to fraction regenerated, independent of spatial identity of fin rays, consistent with previous results in caudal fins^11^ (n=64). C) Average ERK activity over time shows a trend for posterior rays to exhibit lower average activity (n=6, error bars = SEM). D) Maximum projection images of representative fin rays expressing a Wnt activity reporter (*6xTcf/Lef-miniP:dGFP*) in anterior and posterior rays from 3-7 dpa show spatially differential levels. Dotted lines represent area quantified over, where regions outside the width of bony rays are included before maturation of regenerated osteoblasts has begun to clearly occur. E) Total fluorescence dynamics of Wnt activity reporter (in arbitrary units) for anterior and posterior rays of 5 fish. F) Normalized activity level of Wnt reporter within the anterior and posterior rays over time shows the posterior rays peak in expression compared with the anterior at 4 dpa, while anterior ray expression peaks later relative to the posterior rays at 6 dpa (n=5, error bars = SEM). G) Maximum projection images of representative anterior and posterior fin rays showing *fgf20a-*expressing basal epithelial cells between 3 and 10 dpa. Dotted lines represent area quantified over, allowing some addition of cells outside the width of the bony rays before well-defined bone maturation has occurred. H) Total number of cells with *fgf20a* regulatory sequence activation over time in anterior and posterior rays of 6 fish shows an early peak in the number of transcribing cells, which declines as regeneration progresses. I) Normalized number of *fgf20a* expressing cells between anterior and posterior shows no distinct dynamics between anterior and posterior (n=6, error bars = SEM).

Orthogonally, Wnt pathway activity at the distal tip of regenerating caudal fin rays has been shown to be essential for blastemal cell proliferation in early regeneration^37^. To understand its dynamic role during pectoral fin regeneration, we imaged Wnt activity dynamics as they are expressed primarily in mesenchymal cells using an available reporter of downstream (*6xTcf/Lef-miniP:dGFP*)^38^ (Fig. 3D). Across a cohort of 5 fish, we found anteroposterior asymmetries in peak expression levels of this reporter in fibroblast tissue at the distal tip of the growing ray (Figs. 3E,F). Specifically, we found normalized expression levels peaked earlier in posterior fin rays at 4 dpa (0.81 ± 0.08, SEM) compared with anterior (0.51 ± 0.20, SEM), while normalized anterior expression peaked later at 6 dpa (0.92 ± 0.06, SEM) compared to the waning posterior rays (0.59 ± 0.09, SEM). Importantly, these respective peaks between posterior and anterior rays, at around 4 and 6 dpa, match the estimated growth rate times from earlier analyses, suggesting that Wnt activity may be involved or affected by the asymmetry in growth rates during regeneration.

Expression of *fgf20a* in basal epithelial cells is a well-known early signaling event needed for regeneration^39,40^. More recently, it has been argued that the number of expressing cells at the stage of early outgrowth (3 dpa) are involved in instructing the length of tissue eventually regenerated^11^. To understand if differential *fgf20a* transcription might explain the observed Erk and Wnt asymmetry, we quantified the dynamics of *fgf20a* regulatory sequence activation within the regenerating pectoral fin (Fig. 3G). To do this, we live-imaged zebrafish containing both the epithelial-specific promoter *krtt1c19e:H2A-mCherry*, along with the enhancer trap *fgf20a:EGFP* lines^11,39,41^. Across a cohort of 6 longitudinally tracked fish, we indeed found, on average, a higher number of expressing basal epithelial cells in the anterior (4.1 x 10^3^ ± 4.3 x 10^2^ cells, SEM) compared to the posterior (2.7 x 10^3^ ± 3.4 x 10^2^ cells, SEM) at 5 dpa, when both regions possessed their average maximal number of expressing cells (Fig. 3H). Importantly, however, after normalization, we observed no difference between anterior and posterior dynamics, suggesting *fgf20a* gene activation does not exhibit regional asymmetry throughout pectoral fin regeneration (Fig. 3I).

### Bulk RNA-seq analysis identifies cohort of differentially regulated spatial gene patterns

In addition to identifying pathways whose expressions may guide regenerative outgrowth, we also sought to identify genes with anteroposterior asymmetries throughout regeneration. To better account for rarely expressed ligands and other genes that may be exported into extracellular space and therefore missed by single cell methods, we performed bulk RNA-seq across a series of timepoints throughout regeneration (0-12 dpa), splitting the fin into anterior and posterior regions by dividing the tissue between rays 3 and 4 after amputating about half of both pectoral fins of ∼150 female zebrafish and collecting the regenerate (Fig. 4A).

**Figure 4:**
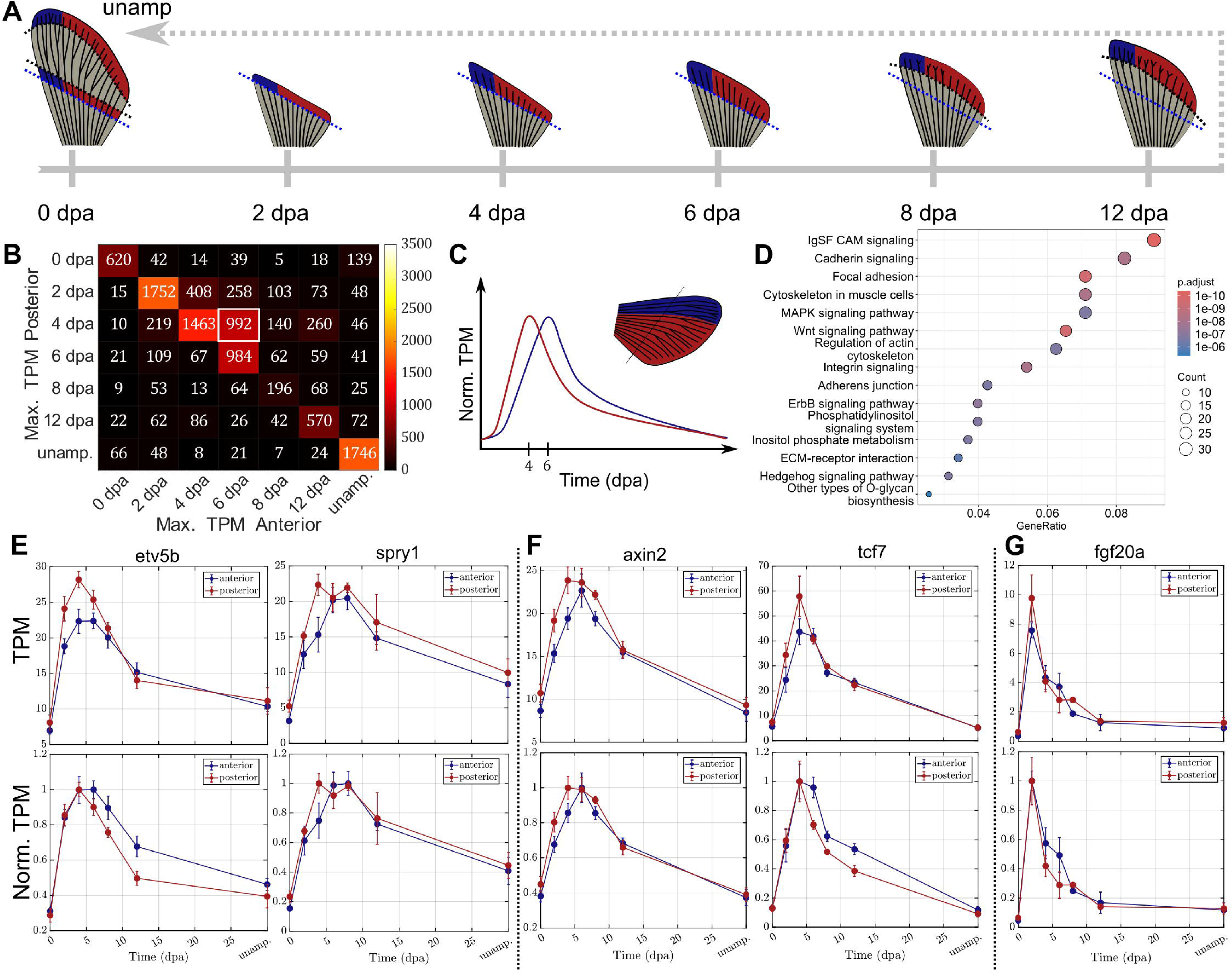
Bulk RNA-seq identifies large subset of genes with anteroposterior asymmetries in expression dynamics. A) Timeline schematic for collection of anterior and posterior tissue across regenerative time and three biological replicates at unamputated, 0, 2, 4, 6, 8, and 12 dpa levels for bulk RNA-seq analyses. B) Heatmap of maximum posterior and anterior timepoints of all genes with TPM expression levels over a threshold of 5 TPM in a majority of timepoints (n=11,235). Of these, most (7,335) reached their maximum expression at the same time independent of anteroposterior identity. However, a large cohort (992) had an earlier maximum expression at 4 dpa in the posterior and 6 dpa in the anterior. C) Schematic representation of normalized TPM over time of an asymmetrically expressed gene. Here, we see over time, from a low expression level at 0 dpa, there is a rapid increase and decrease across regeneration timepoints in both regions of the fin, but there is also a clear spatial difference with respect to which the expression patterns peak. D) KEGG pathway analysis on this cohort of these 992 asymmetric genes shows enrichment of MAPK and Wnt signaling pathways, again suggesting differential expression of genes in these pathways may be responsible for the observed differential dynamics in these regions. In addition, mechanical signaling pathways were also found to be highly enriched, suggesting the importance of these pathways in guiding regeneration. E) Total and normalized TPM plots of downstream targets of Erk signaling with asymmetric expression profiles include the transcription factor *etv5b* and MAPK inhibitor *spry1*. F) Total and normalized TPM plots of downstream targets of Wnt signaling with asymmetric expression profiles include the β-catenin stabilizer *axin2* and transcription factor *tcf7*. G) Total and normalized TPM plots of *fgf20a* expression do not show spatiotemporally distinct behaviors between anterior and posterior regions of the fin.

After normalization through converting raw RNA counts to transcripts per million (TPM) and thresholding for genes with TPM > 5 of at least 5 timepoints in both regions of the fin, we identified the time of maximum gene expression levels in both the anterior and posterior sections of the fin. Of the 11,235 genes that satisfied these requirements, we found most genes (7,331) reach their maximum levels at the same time independent of A/P identity (Fig. 4B). However, we also found a large population of genes that peak in the posterior at 4 dpa and 6 dpa in the anterior (992), consistent with our maximum growth rate data (Fig. 4C). Pathway analysis on these genes again highlighted important mechanical signaling pathways, including cell adhesion, cadherin signaling, cytoskeleton regulation, and extracellular matrix (ECM)-receptor interactions, in addition to the MAPK, Wnt, and Hedgehog pathways (Fig. 4D). Exploring expression dynamics of downstream targets of the Erk and Wnt pathways identified example genes with asymmetries in the peak of anterior expression, as well as ones where posterior expression levels decay more slowly than anterior (Figs. 4E, 4F). In contrast, the expression dynamics of *fgf20a* were shown to behave symmetrically between the anterior and posterior regions of the fin (Fig. 4G). Together, these bulk RNA-seq analyses on expression dynamics provide additional evidence to support the observed spatial asymmetry with respect to Wnt and Erk activity during early regeneration and suggest further exploration of mechanochemical pathways for future studies to study their role and effects on appendage regeneration.

## Discussion

Appendage regeneration is a complex process involving cooperation and communication across cell types to coordinate cell proliferation and migration to replace missing tissue^42^. While transcriptomic analyses can provide insights into existing signaling relationships, live, longitudinal imaging of cell- and tissue-level dynamics provide a full, dynamic. In this work, we introduce new tools that enable *in vivo* longitudinal imaging of pectoral fin regeneration. We additionally expect modifications of these methods and of our 3D printed device will enable longitudinal imaging for limb regeneration studies in other animals. Future improvements on this device should seek to increase the time the animal can remain immobilized or enable breathing without disrupting image acquisition of the pectoral fin. Such modifications should enable extended studies to late times, in addition to allowing other quantitative imaging protocols, such as photoconversion, to become possible in this system.

Our results reveal a key property that has not been characterized in other regenerating appendages: the existence of distinct growth dynamics with respect to anteroposterior position and not simply due to differences in length amputated. Analyzing the relationship between gene expression in regenerative cell types and estimated growth rates implicated the MAPK and Wnt signaling pathways, consistent with previous works into zebrafish appendage regeneration studies^11,32–34^. Furthermore, confocal imaging revealed that while early Fgf expression dynamics were spatially consistent, there exist apparent A/P asymmetries in Erk and Wnt activity dynamics. Further transcriptomic analyses comparing A/P regions across regeneration timepoints identified a large group of genes whose maximum expression is reached earlier in the posterior than anterior enriched for genes in the MAPK and Wnt signaling pathway, as well as many pathways related to cell mechanics and mechanotransduction.

One key novelty of these results lies in the spatiotemporal asymmetry of Wnt activity across the regenerating tissue. Previous work characterizing and perturbing Wnt activity during regeneration has largely been focused at fixed timepoints during blastema formation or early outgrowth^31,34,36^. On the other hand, how Wnt activity dynamics affect fin ray growth and cell proliferation during later stages of outgrowth has been less well-studied. In particular, the signaling or morphological events that may drive this asymmetry in Wnt signaling is an interesting direction for future research. As an example, the Hedgehog signaling pathway is also enriched earlier in the posterior region based on our collected bulk RNA-seq data. Interestingly, expression of this pathway has strong anteroposterior asymmetries that arise as early as limb bud development, where Sonic hedgehog (Shh) is localized to the posterior portion of the fin bud and surrounded by a domain of the transcription factor *hand2*, a persistent posterior forelimb marker^9,43,44^. Due to the existence of feedback interactions between the Shh and Wnt signaling pathways in other regenerative contexts, future work will study if and how Shh signaling during development and regeneration may contribute to the onset of Wnt asymmetry within the regenerating pectoral fins^37,45,46^.

The coordination of multiple cell types to generate proper cell proliferation and migration as the tissue grows implies a role for intra- and intercellular mechanics throughout the process. Thus, the effects of mechanics and mechanotransduction as an essential part of regeneration cannot be ignored. Recent work has identified that stiffness of the ECM itself can dictate the difference between regenerative environments and fibrotic scarring in mouse digit tips^47^. Other work has shown feedback between the mechanosensitive protein *yap1* and the Wnt pathway may also guide cell and tissue movements during regeneration, providing a possible avenue to explore how mechanochemical coupling may generate the observed asymmetric dynamics of regenerating pectoral fins^48–51^. Overall, studying and quantifying mechanical properties such as cell density, tissue stiffness, and the effects of perturbing these characteristics within regenerating fin rays will require *in vivo* biosensors and mechanical tools, yielding a strong direction for future research into how tissues are re-formed and re-built after injury.

## Methods

### Zebrafish husbandry and pectoral fin amputations

Wildtype and transgenic fish derived from Ekkwill strains ranging from 4 to 16 months in age were used throughout all studies. Due to the known defects in pectoral fin regeneration of males, only females were considered in these analyses. Fish were housed between 4 and 7 per liter in Pentair Aquatic Habitat Recirculating Multi-Rack Systems and fed 2-3 times per day. Water temperature was kept between 26 and 28.5°C and the fish were kept on a 14:10 hour light:dark cycle. To perform pectoral fin amputations, fish were anesthetized in 0.75% v/v 2-phenoxyethanol (Sigma-Aldrich) in fish water until breathing slowed, then placed upside down within the imaging chamber, where the right fin was cut with iridectomy scissors, as previously^15^. All experiments using zebrafish were approved by the Institutional Animal Care and Use Committee at Duke University. Transgenic lines used in this study include *Tg(osx:H2A-mCherry)^pd310(20^)*, *Tg(osx:ErkKTR-mCerulean)^pd2001(33)^*, *Tg(6xTcf/Lef1-miniP:dGFP)^isi04(38)^, *Tg(krtt1c19e:H2A-mCherry)^pd309(41)^, fgf20a:EGFP(39)**, and the *longfin (lof)^d2^* mutant(^21^).

### Microscopy

All confocal images were acquired on a Leica Stellaris confocal microscope using LAS X software with a 25x FLUOTAR water immersion lens (Leica 15506374). Full images were acquired at 0.75x zoom, resulting in 1.21-micron x 1.21-micron x 1.21-micron voxel size to ensure best downstream image processing capabilities. For acquisition, zebrafish were anesthetized in 4 mL 0.01% tricaine (MS-222, VWR) in 100 mL of fish water until swimming ceased and gilling slowed sufficiently. Fish were then placed into the imaging mold upside down and the right (cut) pectoral fin was splayed out. Once properly arranged, a small amount of cooling 1% agarose was placed on the fin tissue until immobilized.

The imaging apparatus, which consisted of the 3D-printed mold within a previously described imaging chamber attached with modeling clay, was filled until submerged with a solution of 6 mL 0.01% tricaine (MS-222, VWR) in 200 mL fish water, fully covering the fish for imaging^11,20^. Gill movements were monitored throughout imaging, and, if stopped, the fish was removed from the system to recover before any further image acquisition. After acquisition finished, fish were immediately returned to fish water and recovered. During longitudinal imaging studies, fish were returned to the aquatic system between time points.

All low-magnification images were acquired on a Zeiss AxioZoom V16 using ZEN Pro software. Here, fish were anesthetized in 0.075% v/v 2-phenoxyethanol in fish water and placed within a pectoral fin imaging mold upside down inside a petri dish. Fish were returned to fish water after image acquisition.

### Data analysis

#### Ray length measurements

Length amputated was measured from low-magnification, non-compressed TIFF files of pectoral fins before and after amputation in FIJI. Here, each fin ray was measured from the base where it connects with the distal radial to the amputation plane stump or distal fin ray tip. Length amputated was defined as the difference in length between the unamputated ray and amputation stump measurements. Regenerated lengths were calculated as the difference in length between the distance from the distal radial to distal fin ray tip and the post amputation length at each timepoint. For confocal images, regenerated length was measured manually and defined as the distance between the identified amputation plane and the distal tip of visible tissue. In all cases, fraction regenerated was defined as the ratio between regenerated length to amputated length.

#### Preprocessing of confocal images

When necessary, acquired z-stacks tiling each regenerating ray were stitched into a single 3D image using previously published custom software implementing cross-correlation^33^. These z-stacks were then rotated to minimize the z-coordinate and oriented with the amputation plane on the left-hand side of the regenerating ray. Because osteoblasts consist of two layers of hemi-rays, only the layer closest to the imaging plane was considered. Z-stacks were further equalized for nuclear segmentation, as previously described^11,33^. For further downstream quantifications, such as for expression measurements, non-equalized z-stacks were used. In all cases, codes used in previous analyses based on Tracking with Gaussian Mixture Models (TGMM) software were implemented^52^.

#### ERK activity quantification

Erk activity measurements were performed as in previous studies^11,33^. Briefly, nuclear segmentation was performed, and each nucleus mask was dilated and subtracted from the initial mask to create a cytoplasmic shell. The Erk-KTR levels were quantified as average/pixel in the cytoplasmic shell divided by the average/pixel in the nuclear mask. The Erk activity was taken to be this ratio minus 0.8, which was done as in previous work, based on the Erk activity observed in cells treated with an inhibitor acting directly upstream of Erk activation^11^. The average value across all nuclei in each sample was taken to be the Erk activity of the fin ray at each timepoint.

#### Quantification of Wnt activity

To quantify Wnt activity throughout the regenerating fin ray, fish expressing the marker *6xTcf/Lef-miniP:dGFP* were used, and expression was primarily observed in the mesenchymal cells near the distal tip of regenerating rays^38^. From here, the collected data was run through the TGMM pipeline and objects were identified. The total expression was calculated as the sum of reporter fluorescence throughout the fin ray, and normalizing within the same ray across timepoints generated a relative expression level. Across anterior and posterior fin rays and across samples, the average relative expression level of Wnt activity was recorded, generating a visualization of the asymmetry in Wnt activity.

#### Quantification of fgf20a:EGFP-expressing cells

Counting *fgf20a*:EGFP-expressing cells was done via imaging fin rays from fish containing both *fgf20a*:EGFP and *krtt1c19e*:H2A-mCherry constructs and quantified as performed previously^11^. Images were collected and each fin ray was cropped such that soft tissue surrounding the bony rays was included for the final quantifications. Epithelial nuclei marked by the *krtt1c19e* promoter were segmented with TGMM and these masks were overlaid with the *fgf20a*:EGFP signal, which is homogenously distributed throughout the cell and nucleus. The average expression per pixel within each nucleus was calculated, and all cells with an average expression above 1 arbitrary unit were taken to be *fgf20a*-expressing. Total numbers per fin ray across time were calculated and averaged across samples. To normalize, the number of positive cells for all times within each fin ray sample were normalized by the maximum number of positive cells within the ray to generate a dynamic plot of normalization number of cells within each spatial region over time.

### Single nuclei RNA-seq collection and analysis

Adult female zebrafish pectoral fins on both sides were amputated at the 50% level and collected initially to serve as an unamputated control. For further 2, 4, and 7 dpa timepoints, an additional amputation was performed approximately one bone segment proximal to the initial amputation plane. For each collected sample, the removed portion was sectioned to include the following portions: rays 1 and 2 (AP1), rays 3 and 4 (AP2), rays 5 and 6 (AP3), rays 7 and 8 (AP4), and the remaining rays 9 to the posterior end of the fin (AP5). Each collected timepoint consisted of fin tissue from 25 females (50 total fins), sectioned into the five portions. For each sample, nuclei isolation was performed based on previously established protocols^53^. Briefly, for each sample, tissue was flash frozen in liquid nitrogen and maintained at −80° C until sample preparation. These frozen tissues were transferred to a cooled Dounce homogenizer (Sigma) with 1 mL of Homogenization buffer containing 0.2 U/ µL RNAse inhibitor (ThermoFisher) and 1:200 PIC III (Sigma). Samples were manually homogenized at least 10 times each with the A and B Pellet Pestle and incubated on ice. Samples were then filtered through a 40-micron cell strainer and centrifuged in a swinging bucket rotor at 300 g for 10 minutes at 4° C. Pellets were then resuspended in PBS with 2% BSA and 1 U/μL RNAse inhibitor. Nuclei were counted on a hemocytometer. Sample preparation, including nuclei fixation, barcoding, cDNA amplification, and library preps were prepared via provided manufacturer instructions using the Parse Biosciences Evercode Whole Transcriptome kit and sequenced through BGI-Biosciences/Innomics to yield 150 bp paired reads. Raw reads were mapped to the *Danio rerio* reference genome GRCz11. FASTQ files were processed with Trailmaker (Parse Biosciences).

Downstream analyses were performed in R (v4.6.1) using Seurat (v5.5.1)^54^, DoubletFinder (v2.0.6)^55^, CELDA (v1.28.0)^56^, harmony (v2.0.5)^57^, and UCell (v2.16.0)^58^. As a first pass, nuclei with exceptionally low total genes detected (<300) or with greater than 5% mitochondrial transcript content were removed, along with any genes detected in only a few nuclei (<20). For each sample collected, potential doublets were identified with DoubletFinder and removed during early QC^55^. Raw counts matrices were updated through estimated contribution of contamination from single nuclei collection process, as calculated through the decontamination pipeline within CELDA, and cells with estimated >40% contamination scores were removed from further analyses^56^. Count matrices were log-normalized in Seurat, the most highly variable genes were identified, and principal components analysis (PCA) was performed on this data^54^. Examination of an elbow plot for the variance of each principal component was consistent with choosing the first 30 principal components for further analyses. Batch correction was performed with Harmony on the reduced PCA dimensions, and Uniform Manifold Approximation and Projection (UMAP) was performed. Unsupervised clustering was performed with FindClusters for all resolutions between 0.1 and 1.2 in 0.1 increments, and a final resolution of 0.6 was determined by inspection based on the number of clusters generated. Cell types were determined through a combination of identifying the most highly differentially expressed genes per cluster, in addition to the gene expression patterns and cluster identities of canonical gene markers taken from previous works and datasets^23–27,58^. Based on this cell type identification, determined mesenchymal and basal epithelial cells were isolated and extracted into their own dataset for pseudobulk analyses.

Transcript counts for each mesenchymal and basal epithelial cell were aggregated by time point collected and by anteroposterior position AP1-AP5 with the AggregateExpression function in Seurat^54^. To generate estimated counts per million (CPM), these aggregated sums were normalized by the number of cells in each cohort. For each position AP1-AP4, previous experiments performed allowed for an estimated growth rate of regeneration to be determined across all timepoints. Thus, for each gene, plotting estimated growth rate against aggregated CPM across AP1-4 and 2, 4, and 7 dpa could yield a corresponding Spearman correlation coefficient. Allowing for a time delay from gene expression to further tissue growth, the Spearman correlation coefficient was determined for all time delays between 0 and 4 days in 0.1-day increments. The maximum Spearman correlation (in absolute value), along with the corresponding time delay associated with this relationship was recorded. All genes with an absolute value of the Spearman correlation above 0.7 and with CPM above 1 for at least one datapoint were considered significant. Positively and negatively associated genes were kept for both basal epithelial and mesenchymal cell populations for each of the two regeneration dynamic experiments. Within each cell type, genes that overlapped across both experiments were kept for further pathway enrichment analysis studies using clusterProfiler (v4.20.0)^59^. Using KEGG annotations, enrichment analyses were performed on cell lists and the top 15 pathway terms by adjusted p-value were displayed^30^.

### Bulk RNA-seq collection and analysis

To extract and compare gene expression profiles between anterior and posterior portions of regenerating pectoral fin rays across timepoints, the most distal ∼500 microns from newly amputated fin rays was collected as an unamputated control, as well as the ∼500-micron portion at the amputation plane as a negative, 0 dpa control. Additional amputations collected the full regenerate from 2 dpa, 4 dpa, and 6 dpa, as well as the most distal ∼500-micron portion of the regenerated fin at 8 dpa and 12 dpa. In all cases the collected tissue was separated between the third and fourth fin ray to separate into anterior and posterior portions. For each cohort, approximately 10 female fish (20 fins) were amputated for each of three biological replicates, yielding a total of ∼150 fish. After collection, all tissue samples were placed in a 2.0 mL RNase-free Safe-Lock tube (Eppendorf) with Tri-Reagent (Sigma) and a 5 mm stainless steel bead (Qiagen), before homogenizing in a TissueLyser (Qiagen) for 20 seconds at 30 Hz. From here, RNA was extracted with chloroform and precipitated with isopropanol and carrier glycogen. DNA was digested with added DNase, and the solution was purified via an RNA Clean & Concentrator Kit (Zymo). Library preparation and sequencing was performed by Innomics, Inc.

RNA-seq reads were mapped by STAR (v2.7.11)^60^ and normalized by gene length using the GRCz11 zebrafish genome annotations. Raw counts by gene length were additionally normalized across samples to generate transcripts per million (TPM) for downstream analyses. To identify which genes had asymmetric A/P expression profiles throughout regeneration, TPM values were first averaged across biological replicates. Next, this average TPM in both anterior and posterior samples was checked against a threshold (5 TPM) across a majority (5) of the collected timepoints. Each gene was then characterized based on what timepoint its maximum expression level was reached in the anterior and posterior to yield a heatmap based on expression level. Genes whose maximum levels were reached at 4 dpa in the posterior and 6 dpa in the anterior were collected and pathway enrichment analysis using clusterProfiler (v4.20.0) was performed^59^. Here, KEGG enrichment analyses were also performed, with the top 15 pathway terms by adjusted p-value again being shown^30^.

## Data availability

The snRNA-seq and bulk RNA-seq datasets generated in this work will be submitted to online databases to be available upon publication. In addition, the .stl file to generate the 3D-printable mold, specifical biological materials used in this work, as well as codes written for downstream analyses of collected data and modeling are available upon request.

## Supplemental Figures

**Figure S1:**
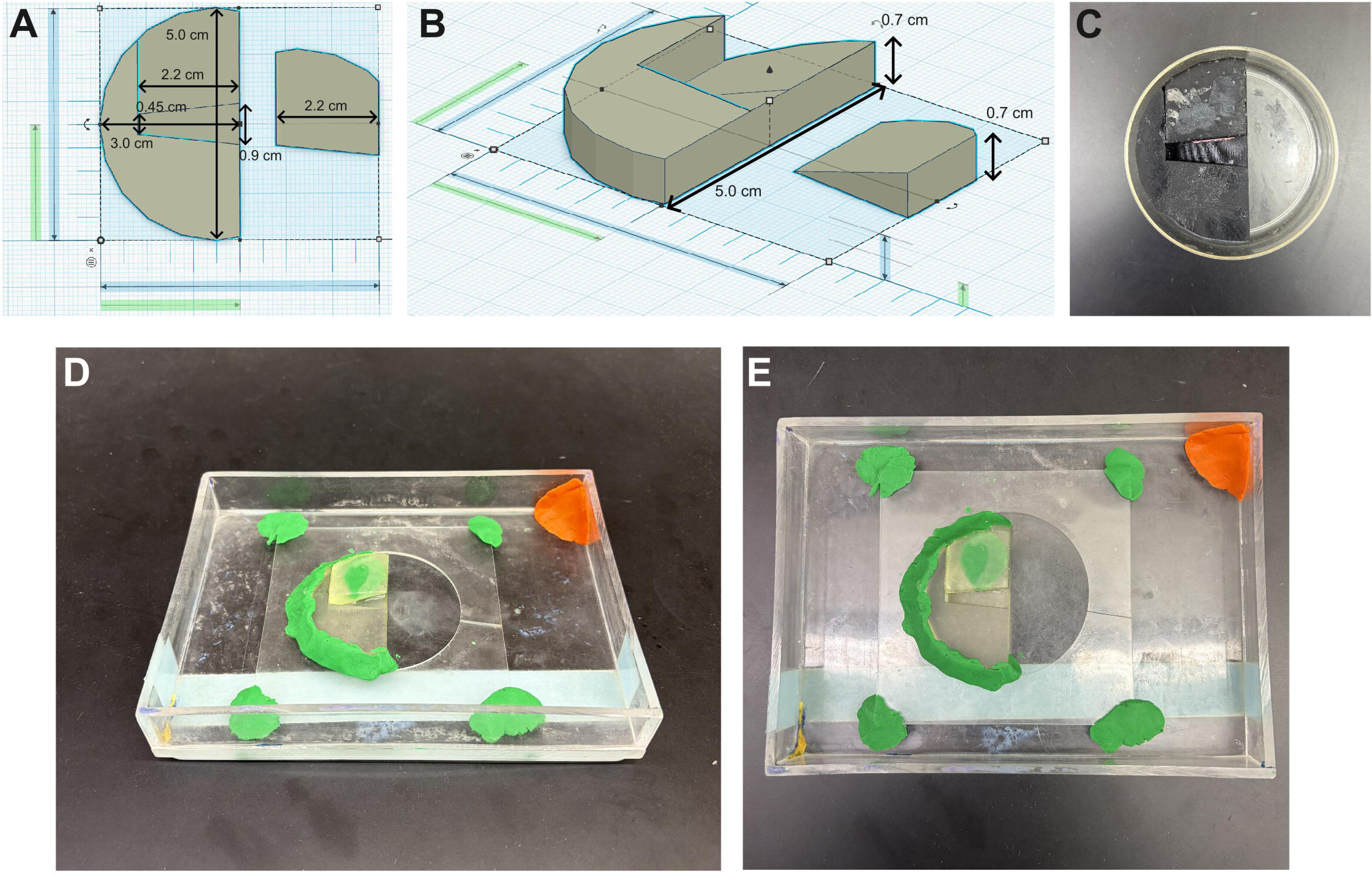
3D printable mold for pectoral fin imaging and imaging setup. A) Top-down view and B) transverse view built in TinkerCAD of device for pectoral fin imaging. This apparatus consists of a roughly semi-circular cylinder with diameter 5.0 cm and height 0.7 cm with a small piece 2.2 cm long cut from it. This piece is also 0.7 cm tall on one side, with slope decreasing down until it reaches 0 on the opposite side, allowing it to be wedged into the imaging device. When fit to meet the circular edges, there is a small crevice for immobilizing fish that is 0.45 cm on one side and 0.9 cm on the opposite side, though this can be adjusted to fit the width of any fish. C) 3D-printed mold inside a small Petri dish to be used for dissection scope imaging. The insertable piece is attached to the rest of the device via modeling clay. D) Transverse and E) top-down view of 3D-printed mold for confocal imaging positioned and mounted inside a plexiglass dish previously used for scale regeneration and caudal fin regeneration imaging^11,20,33^, manufactured by the Duke Machine Shop. The length of the internal plate with circle removed is 9.0 cm, with a diameter of 6.0 cm and height 0.1 cm. The thickness of the bottom of the dish is 0.8 cm, the thickness of all side walls is 0.3 cm, and the inner length, width, and height of the chamber is 15.4, 11.0, and 2.0 cm, respectively.

**Figure S2:**
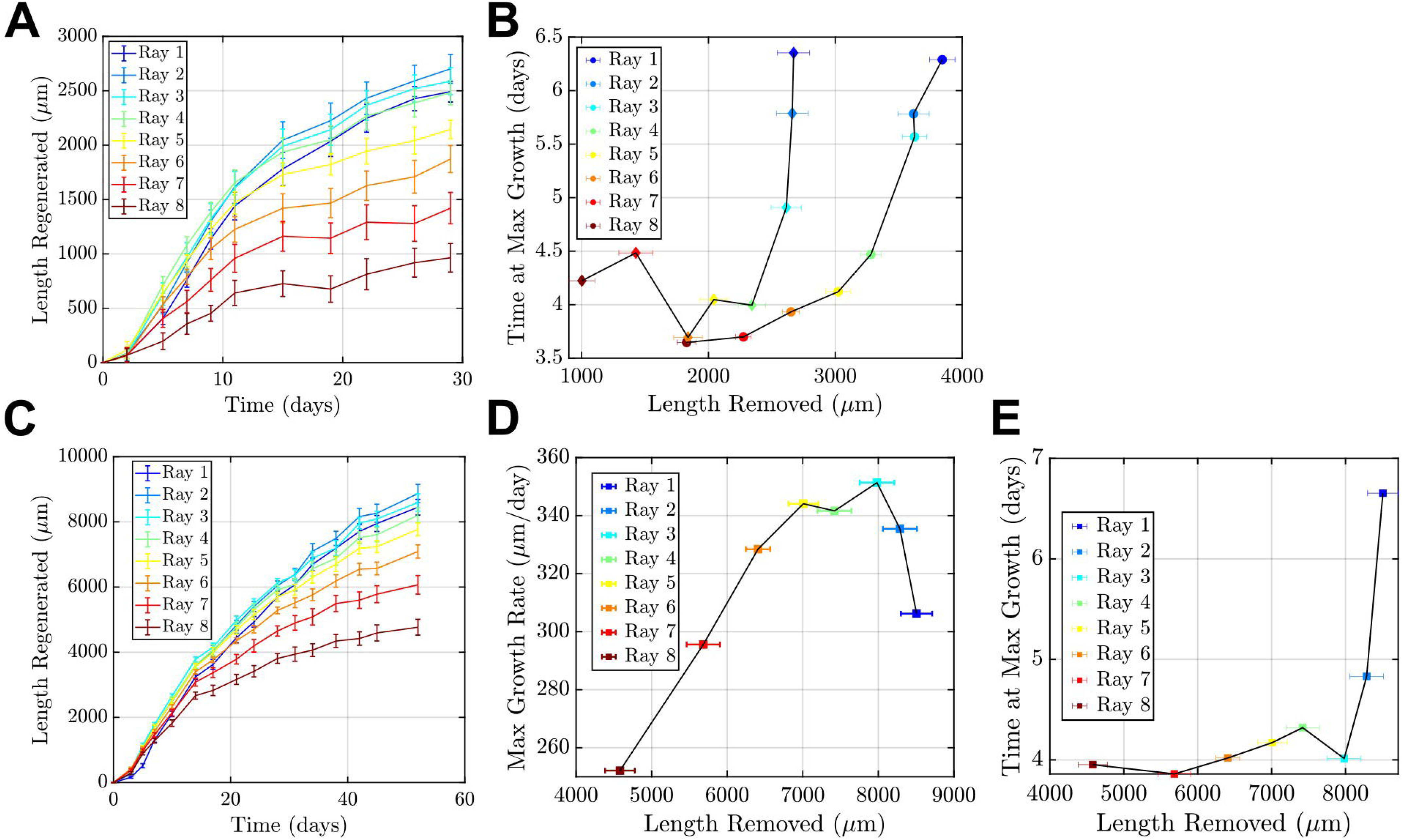
Pectoral fin ray dynamics, maximum growth rate, and time to reach maximum growth for shorter amputation and *longfin*. A) Plot of average total length regenerated as a function of time (n=11) for the less severe, proximal amputation (∼25% removal) experiment. Rays 1 (most anterior, dark blue) through 8 (most posterior, dark red). Across fish, the most anterior rays’ dynamics (1-3) again appeared qualitatively distinct from the posterior rays’ (4-8), prompting further analyses (error bar = SEM). B) Plot of estimated time at maximum growth rate as a function of length removed for the both the proximal (n=10) and distal (n=11) amputation experiments. Here, a clear distinction between the peak growth rate of the posterior (4-8) and anterior rays (1-3) can be observed, independent of length amputated (error bar = SEM). C) Plot of average total length regenerated as a function of time (n=12) for ∼50% amputation experiment in *longfin* mutant fish. Rays 1 (most anterior, dark blue) through 8 (most posterior, dark red). Across fish, as in the wildtype experiments, the most anterior rays’ dynamics (1-3) again appeared qualitatively distinct from the posterior rays’ (4-8) (error bar = SEM). D) Plot of estimated maximum growth rate as a function of length removed for the *longfin* fish (n=12). We again note the apparent linear relationship between maximum growth rate and length removed for posterior rays, while anterior rays do not fit this relationship as well (error bar = SEM). E) Plot of estimated time at maximum growth rate as a function of length removed for the *longfin* (n=12) amputation experiment again show a deviation, more dramatically for Rays 1 and 2, of anterior rays with respect to when the estimated maximum growth rate is reached (error bar = SEM).

## Acknowledgments

The authors thank A. Rich and X. Lu for helpful discussions, N. Joshi and T. Whitlow for troubleshooting assistance, members of the Di Talia and Poss labs for feedback, and staff members of the Duke Zebrafish Facility. R.D. was funded via support from the Jane Coffin Childs Memorial Fund for Medical Research and the Duke Regeneration Center. S.B. was funded by support from the Duke Center for Advanced Genome Technologies and the National Institutes of Health via grant K99 HD121969. S.D.T was funded by an award in Innovative Stem Cell Science from the Shipley Foundation, Inc. S.D.T and K.D.P. were funded by the National Institutes of Health (NIH) via grants R01 AR076342 (S.D.T., K.D.P) and R01 HD105033 (K.D.P.).

